# The interplay between Mobilome and Resistome in *Staphylococcus aureus*

**DOI:** 10.1101/2024.06.11.598472

**Authors:** Rachel Contarin, Antoine Drapeau, Pauline François, Jean-Yves Madec, Marisa Haenni, Emilie Dordet-Frisoni

**Affiliations:** INTHERES, Université de Toulouse, INRAE, ENVT, Toulouse, France; Anses – Université de Lyon, Unité Antibiorésistance et Virulence Bactériennes, Lyon, France

**Author notes:** Corresponding author: Emilie Dordet-Frisoni,; Marisa Haenni. These authors contributed equally to the work (Marisa Haenni, Emilie Dordet-Frisoni).

## Abstract

Antibiotic resistance genes (ARGs) in *Staphylococcus aureus* can disseminate vertically through successful clones, but also horizontally through the transfer of genes conveyed by mobile genetic elements (MGEs). The underexplored MGE/ARG associations in S*. aureus* favor the emergence of multidrug-resistant clones, posing a significant threat to human and animal health.

This study investigated the interplay between the mobilome, encompassing MGEs, and the resistome, the collection of ARGs, in more than 10,000 *S. aureus* genomes from human and animal sources. The analysis revealed a remarkable diversity of MGEs and ARGs, with plasmids and transposons being the main carriers of resistance genes. Numerous MGE/ARG associations were identified, suggesting that MGEs play a critical role in the dissemination of resistance. A high degree of similarity was observed in MGE/ARG associations between human and animal isolates, highlighting the potential for unrestricted spread of ARGs between hosts. While clonal expansion is a major driver of resistance dissemination in *S. aureus*, our results showed that MGEs and their associated ARGs can spread across different strain types (STs), favoring the evolution of these clones and their adaptation in selective environments. The high variability of MGE/ARG associations within individual STs and their spread across several STs highlight the crucial role of MGEs in shaping the *S. aureus* resistome.

Overall, this study provides valuable insights into the complex interplay between MGEs and ARGs in *S. aureus*, emphasizing the need to elucidate the mechanisms governing the epidemic success of MGEs, particularly those implicated in ARG transfer.

**Importance:** The research presented in this article highlights the crucial importance of understanding the interactions between mobile genetic elements (MGEs) and antibiotic resistance genes (ARGs) carried by *Staphylococcus aureus*, a versatile bacterium that can be both a harmless commensal and a dangerous pathogen for humans and animals. *S. aureus* strains represent a major threat due to their ability to rapidly acquire and disseminate ARGs. By analyzing a large dataset of *S. aureus* genomes, we highlighted the substantial role of MGEs, in particular plasmids and transposons, in the dissemination of ARGs within and between *S. aureus* populations, bypassing the host barrier. Given that multidrug-resistant *S. aureus* strains are classified as a high-prioritypathogenby global health organizations, this knowledge is crucial for understanding the complex dynamics of transmission of antibiotic resistance in this species.

## Introduction

*Staphylococcus aureu*s is a versatile bacterium that can be a commensal microorganism and a lethal pathogen for both humans and animals. The large range of symptoms caused by *S. aureus* in each host is the result of the intricate interplay between complex factors, including host health status, genetic composition of the *S. aureus* strain and the site of infection (1–3). The overuse and misuse of antibiotics in both human healthcare and veterinary medicine have promoted the emergence of numerous antibiotic-resistance genes (ARGs) and the dissemination of multidrug-resistant clones (4). As a result, this species has been classified by the WHO (World Health Organization) as high-priority for the development of new antibiotic targets, and by the WOAH (World Organization for Animal Health) on the surveillance list of multi-resistant pathogens. Studies on antimicrobial resistance (AMR) in *S. aureus* have primarily focused on methicillin-resistance, due to the presence of the *mecA/mecC* genes, which confer resistance to all beta-lactam antibiotics. Consequently, *S. aureus* strains are commonly classified as methicillin-resistant (MRSA) or methicillin-susceptible (MSSA) isolates (5). However, *S. aureus* can acquire multiple additional ARGs conferring resistance to all known antibiotic families, shaping the resistome of each isolate (6). In 2018, a review drew up an inventory of 46 ARGs shared between *S. aureus* isolates of human and animal sources, suggesting potential interspecies transmissions (7).

In contrast to a highly clonal core genome, up to 25% of the *S. aureus* genome is composed of accessory genes, primarily encompassing mobile genetic elements (MGEs) that contribute significantly to the adaptability and survival of these bacteria in various environmental conditions by spreading ARGs mainly occurs through horizontal gene transfer (HGT) (8, 9). MGEs are DNA fragments that can promote intra- or intercellular transfer of genetic material, and play a crucial role in the dissemination of ARGs. The staphylococcal mobilome (set of MGEs) exhibits a remarkable variability (8, 10–13). *S. aureus* MGEs include (i) plasmids (extrachromosomal genetic elements that can be conjugative, mobilizable, or non-mobilizable in *S. aureus*) (12), (ii) integrative and conjugative elements (ICEs) (MGEs that integrate into the bacterial chromosome and have the potential to excise and transfer themselves to other bacteria via conjugation) (14, 15), (iii) insertion sequences (IS) and transposons (Tn) (chromosomally-integrated MGEs that are often associated with other MGEs, in single or multiple copies) (13), (iv) prophages (MGEs usually integrated into the host’s chromosome, since they can repress the phage’s lytic functions) (16), and (v) genomic islands (such as the staphylococcal chromosomal cassettes (SCC*mec*) specific to *mec*-carrying staphylococci) (13).

It is usually admitted that *S. aureus* disseminate through clonal expansion, spreading resistance genes vertically through waves of successful lineages. While the role of MGEs, particularly plasmids, has been extensively studied in Enterobacterales, few studies have addressed the diverse MGEs and associated ARGs carried by *S. aureus* (8, 9, 13, 17). Likewise, no large-scale analyses have been conducted to localize ARGs across *S. aureus* genomes from diverse sources and hosts. To fill in this knowledge gap, this study aimed at (i) identifying the most prevalent MGEs and ARGs in 10,063 *S. aureus* genomes from the NCBI public database, (ii) highlighting MGE/ARG associations and (iii) evaluating the role of MGEs in the emergence and spread of antibiotic resistance in *S. aureus*.

## Results

### Diversity of human and animal *S. aureus* genomes

A total of 9,408 human and 655 animal *S. aureus* genomes were analyzed. Animal *S. aureus* genomes came from 15 different hosts, mainly livestock (n=527, see porcine, bovine, avian, ovine and caprine origins in Table S1). Data originated from all continents, with a significant proportion of *S. aureus* of human source (Hm-SA) isolated from North America and Europe, and of *S. aureus* of animal source (An-SA) isolated from Europe and Asia (Figure 1, Table S1). The core genome of all *S. aureus* strains consisted of 1,716 genes, including 86 genes unique to An-SA and 43 genes unique to Hm-SA. A total of 433 distinct sequence types (STs) were identified, but only 21 were found in >1% of either Hm-SA and/or An-SA were identified. Among the 433 STs, 362 (three present in > 1% of Hm-SA) were unique to Hm-SA, 34 (two present in > 1% of An-SA) to An-SA, and 37 (16 present in >1% of Hm-SA and An-SA) were shared between both groups, highlighting the source-specificity of STs (Figure S1A).

**Figure 1.**
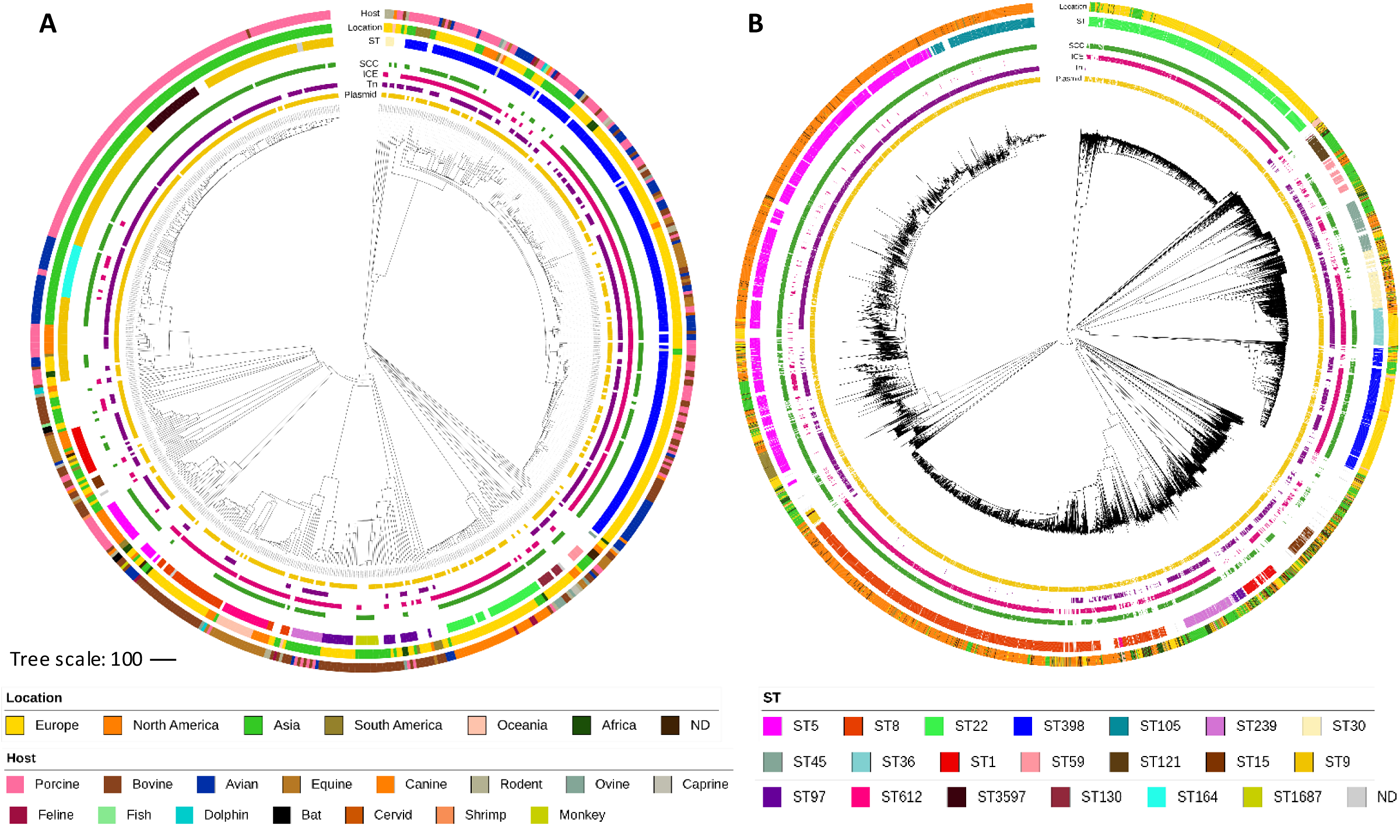
Genomic diversity of *S. aureus* genomes of animal (An-SA) (A) and human (Hm-SA) (B) sources. The phylogenetic trees represent 655 An-SA and 9,408 Hm-SA. This tree based on cgMLST distance was constructed using Neighbor-Joining method and visualized using iTOL v6 (https://itol.embl.de). From outer circle to inner circle, host, location, sequence type (ST), Staphylococcal Chromosomal Cassette (SCC) (green), Integrative and Conjugative Element (ICE) (pink), transposon (Tn) (purple) and plasmid (yellow) were represented. The correspondence between colors and each type of host, location, or ST were indicated below the figure. Only STs with a frequency >1% in Hm-SA and/or in An-SA, were represented. Genomes with no associated ST and/or no location were annotated as “not determined” (ND). As they were present in up to 90% of the analyzed genomes, IS and prophages were not represented.

The most frequently identified STs were (i) the two major livestock associated ST398 (n=216, mainly in Europe) and pig-associated ST9 (n=125, mainly in Asia) (Figures 1A & S1B) in An-SA and (ii) ST5 (n=2,138) and ST8 (n=1,719) in Hm-SA (Figures 1B and S1B).

### Mobile genetic elements arsenal in *S. aureus* genomes

Analysis of accessory genomes highlighted aremarkable diversity of MGEs including ICEs, transposons and plasmids regardless of their human or animal sources (Figure 1, Table 1). A total of 114,845 MGEs, or 59,768, if excluding IS, were detected (Tables 1 & S1), with IS elements, prophages, plasmids and SCC*mec* being the most prevalent, followed by transposons and ICEs (Table 1). Composite transposons (CTn) were the least common MGE detected in *S. aureus* genomes. Transposons and ICEs were found in *S. aureus* genomes irrespective of their origin. In contrast, IS elements and plasmids exhibited greater host specificity, since only 33% and 23% were respectively shared between Hm-SA and An-SA strains (Table 1).

**Table 1.** Prevalence of mobile genetic elements (MGEs) identified in *S. aureus* genomes of animal (An-SA) and human (Hm-SA) origin.

|  | Plasmid |  | Transposon |  | ICE |  | SCC |  | Phage |  | IS |  | CTn |  | All MGEs |  | All MGEs (except IS) |  |
| --- | --- | --- | --- | --- | --- | --- | --- | --- | --- | --- | --- | --- | --- | --- | --- | --- | --- | --- |
| Sources | Hm | An | Hm | An | Hm | An | Hm | An | Hm | An | Hm | An | Hm | An | Hm | An | Hm | An |
| Quantity of MGEs identified | 12,084 | 929 | 6,581 | 499 | 5,287 | 436 | 471 | 7,518 | 867 | 19,124 | 2,873 | 52,204 | 5,585 | 387 | 108,383 | 6,462 | 56,179 | 3,589 |
| Mean of MGEs per strain | 1.5 | 1.7 | 1.4 | 1.2 | 1.1 | 1.2 | 1 | 1 | 2.1 | 1.6 | 5.6 | 4.5 | 1.7 | 1.9 | 11.5 | 9.9 | 6.0 | 5.5 |
| Maximum of MGEs per strain | 8 | 6 | 7 | 3 | 4 | 3 | 1 | 1 | 6 | 5 | 58 | 41 | 17 | 11 | 78 | 57 | 23 | 16 |
| Strains containing MGEs (%tot genomes)* | 8,064 (86%) | 532 (81%) | 4,707 (50%) | 420 (64%) | 4,751 (50%) | 363 (55%) | 7,518 (80%) | 471 (72%) | 9,090 (97%) | 554 (85%) | 9,286 (99%) | 636 (97%) | 3,368 (36%) | 206 (31%) | 9,408 (100%) | 655 (100%) | 9,408 (100%) | 655 (100%) |
| MGEs specific to sources | 203 | 25 | 2 | 0 | 0 | 0 | 12 | 1 | 30 | 4 | 35 | 8 | 10 | 2 | 292 | 40 | 257 | 32 |
| MGEs shared between sources (%MGEs shared)** | 69 (23%) |  | 9 (82%) |  | 3 (100%) |  | 18 (58%) |  | 51 (60%) |  | 21 (33%) |  | 10 (45%) |  | 181 (35%) |  | 160 (36%) |  |

ICE: Integrative and Conjugative Element; SCC: Staphylococcal chromosomal cassette; IS: Insertion Sequence; CTn: Composite transposon.
\* Number of MGE/total number of genomes (in %)
\*\*Number of MGE shared between sources/total number of type of MGEs (in %)

The MGE content varied significantly depending on STs. Plasmids, IS and SCC were present in all major STs (found in >1% of isolates), regardless of their sources. However, specific MGEs could be absent from certain STs, such as transposons that were not identified in ST22 genomes (Figure S2A & B). The mobilome was also diverse among isolates of a same ST, particularly in their plasmid and transposon content, as observed for An-SA ST398 and Hm-SA ST8 (Figure 1).

A wide diversity was also observed for each MGE (Figure 2). Among plasmids, Rep1 was predominant in both An-SA (43%) and Hm-SA (29%) genomes, while others segregated depending on the source, like RepA_N and RepL in Hm-SA (16%) versus Rep_trans in An-SA (24%) (Figure 2). Plasmids also frequently harbored multiple *rep* genes (37% of all Hm-SA plasmids possessing multiple *rep* genesand 23% in An-SA). For transposons, the Tn*554* family was prevalent (30%) in Hm-SA, the Tn*558* family was widespread (27%) in An-SA and the Tn*552* was equally distributed (about 20% in Hm-SA and An-SA) (Figure 2). Concerning ICEs, the ICE*6013* family prevailed in Hm-SA (44% of genomes versus 26% in An-SA), whereas the Tn*916* family exhibited a higher prevalence in An-SA (38%, and 12% for Hm-SA) (Figure 2). For SCC*mec*, type II(2A) and type IVa(2B) were frequent in Hm-SA (27% and 19%, respectively), while type XII(9C2) was the most prevalent (21.5%) in An-SA (Figure 3A). Among prophages, the most common were phi2958PVL and P282 families, present in more than 33% of Hm-SA genomes, while the prophage StauST398 was the most abundant in An-SA (28%) (Figure 3B). For IS, the most common families were IS*Sau6*, IS*Sau3* and IS*1272*, regardless of the source (Figure 3C). IS*Sau4* was very frequent in Hm-SA(41%) and almost absent in An-SA(0.9%) (Figure 3C). Contrarily, IS*256* was widespread in An-SA (36%) but rarely found in Hm-SA (9%) (Figure 3C). Finally, as observed for ISs, the CTn containing IS*Sau6* was the most prevalent, identified in over 23% of *S. aureus* genomes, regardless of their source (Figure 3D). The CTn with IS*Sau4* was abundant in Hm-SA genomes (11.5%) but rare in An-SA genomes (0.5%).

**Figure 2.**
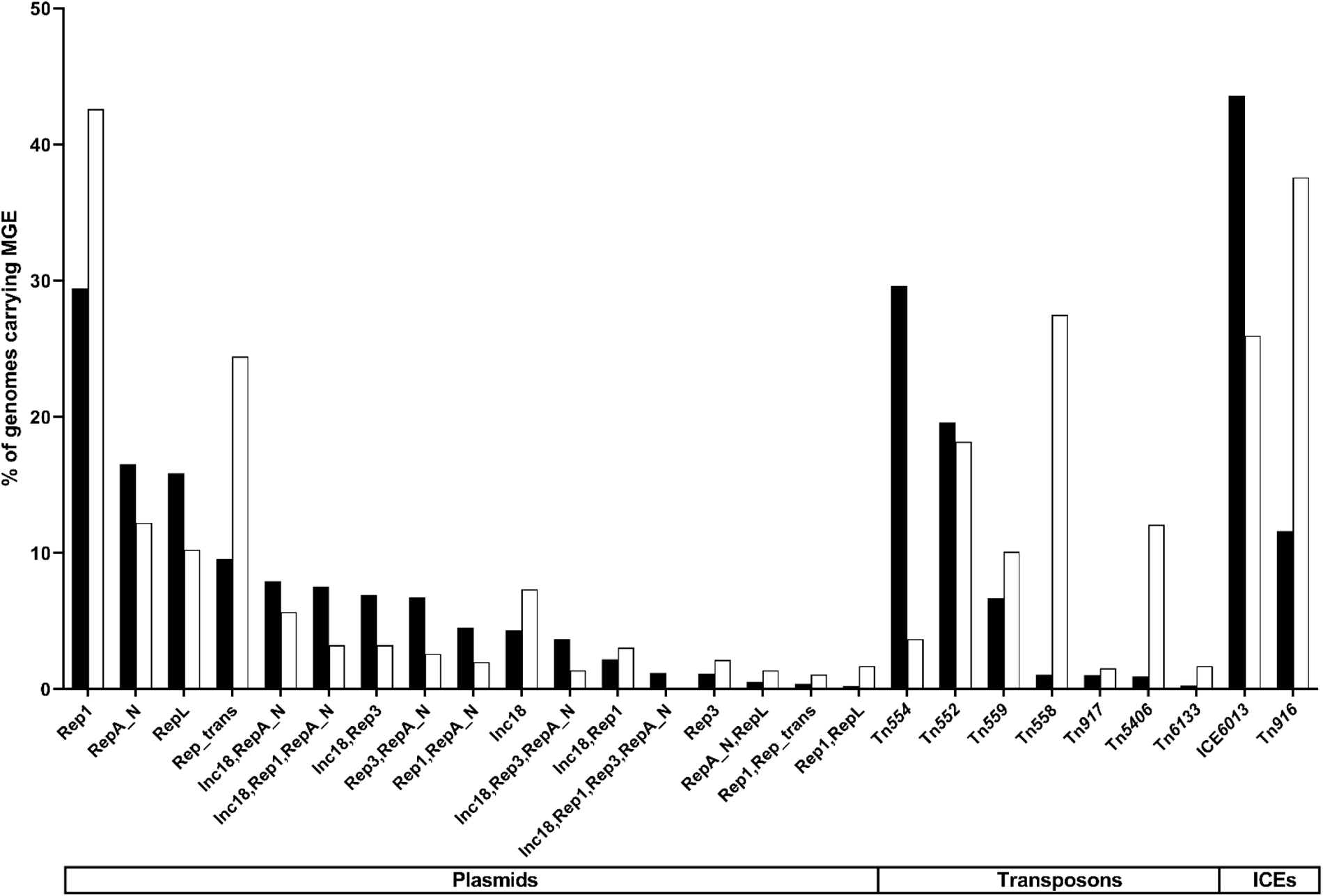
Diversity and occurrence of the most prevalent mobile genetic elements (MGEs) families (plasmids, transposons and integrative and conjugative elements (ICEs)) among *S. aureus* of human (Hm-SA, black) and animal (An-SA, white) origins. Bar chart of the percentage of genomes carrying MGE which corresponds to the number of genome containing at least one MGE out of the total number of genomes analyzed (Hm-SA n=9,408 and An-SA n=655). Only types of MGEs identified in >1% of Hm-SA and/or An-SA genomes are represented in the x-axis.

**Figure 3.**
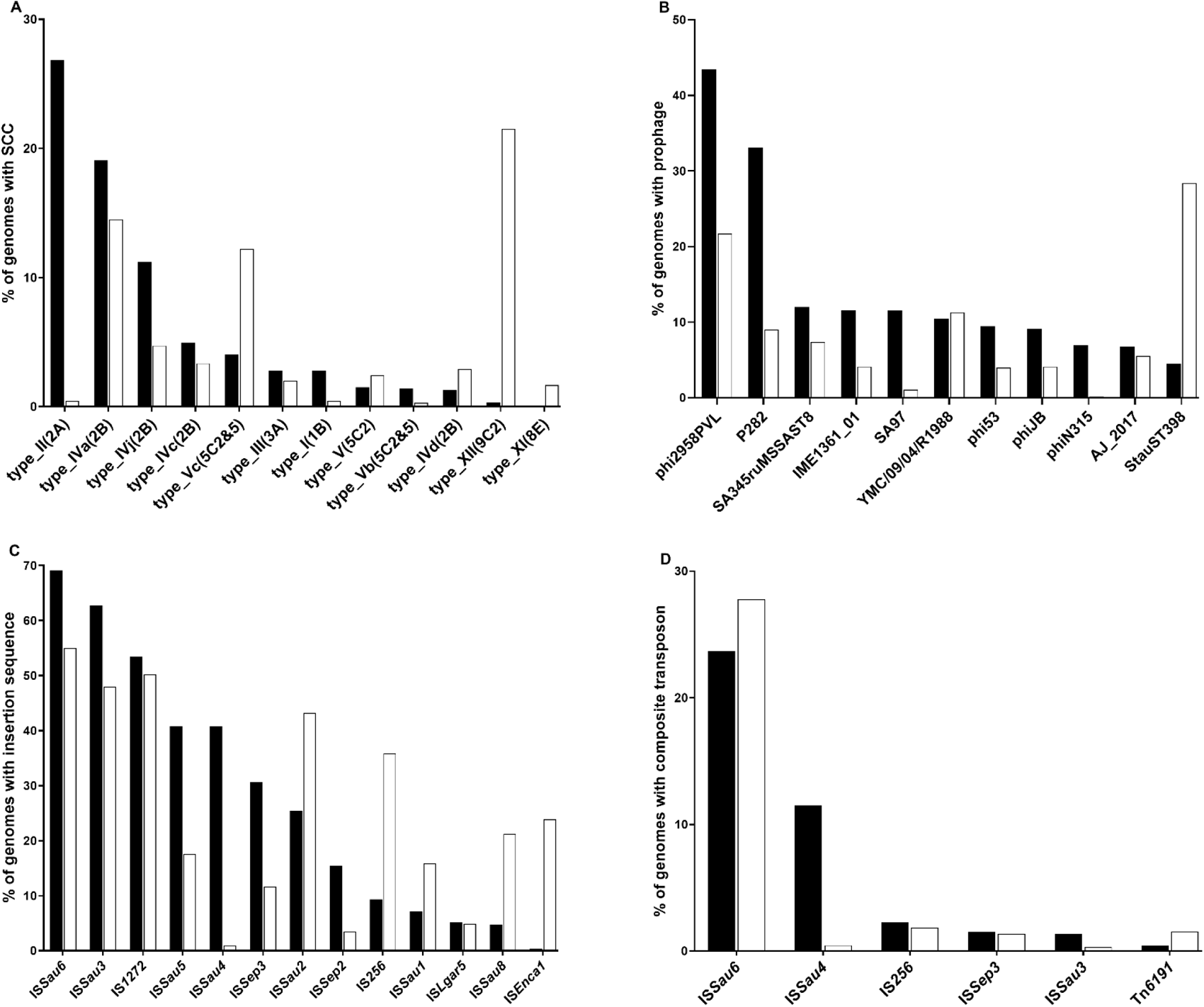
Occurrence and diversity of Staphylococcal Chromosomal Cassette s (SCC) (A), prophages (B), insertion sequence (IS) (C) and composite transposons (CTn) (D) detected families in *S. aureus* of human (Hm-SA, black) and animal (An-SA, white) sources. Bar chart of the percentage of genomes carrying MGE which corresponds to the number of genomes carrying at least one MGE out of the total number of genomes analyzed (Hm-SA n=9,408 and An-SA n=655). For SCC and CTn only MGEs identifiedin >1% of Hm-SA and/or An-SA genomes were represented. For prophages and IS only MGEs identified in >5% of Hm-SA and/or An-SA genomes were depicted.

### *S. aureus* antibiotic resistance genes content

Seventy-eight different ARGs were identified, of which 50 were shared by both Hm-SA and An-SA genomes, 27 were unique to Hm-SA, and one (*optrA*) was unique to An-SA (Figure 4, Table S1). *S. aureus* genomes harbored an average of six or five different ARGs per An-SA or Hm-SA genomes, respectively, with a maximum of 17 per Hm-SA and 15 per An-SA (Table S1). In An-SA isolates, the *lnu*(B) gene was consistently found in tandem with the *lsa*(E) gene (>99%) (Figure S3). Similarly, in Hm-SA some ARGs were frequently collocated(>90%) as *ant*(*9*)*-Ia* and *erm(A)*, *ant*(*6*)*-Ia* and *aph(3’)-III*, and *mph(C)* and *msr(A)* (Figure S3).

**Figure 4.**
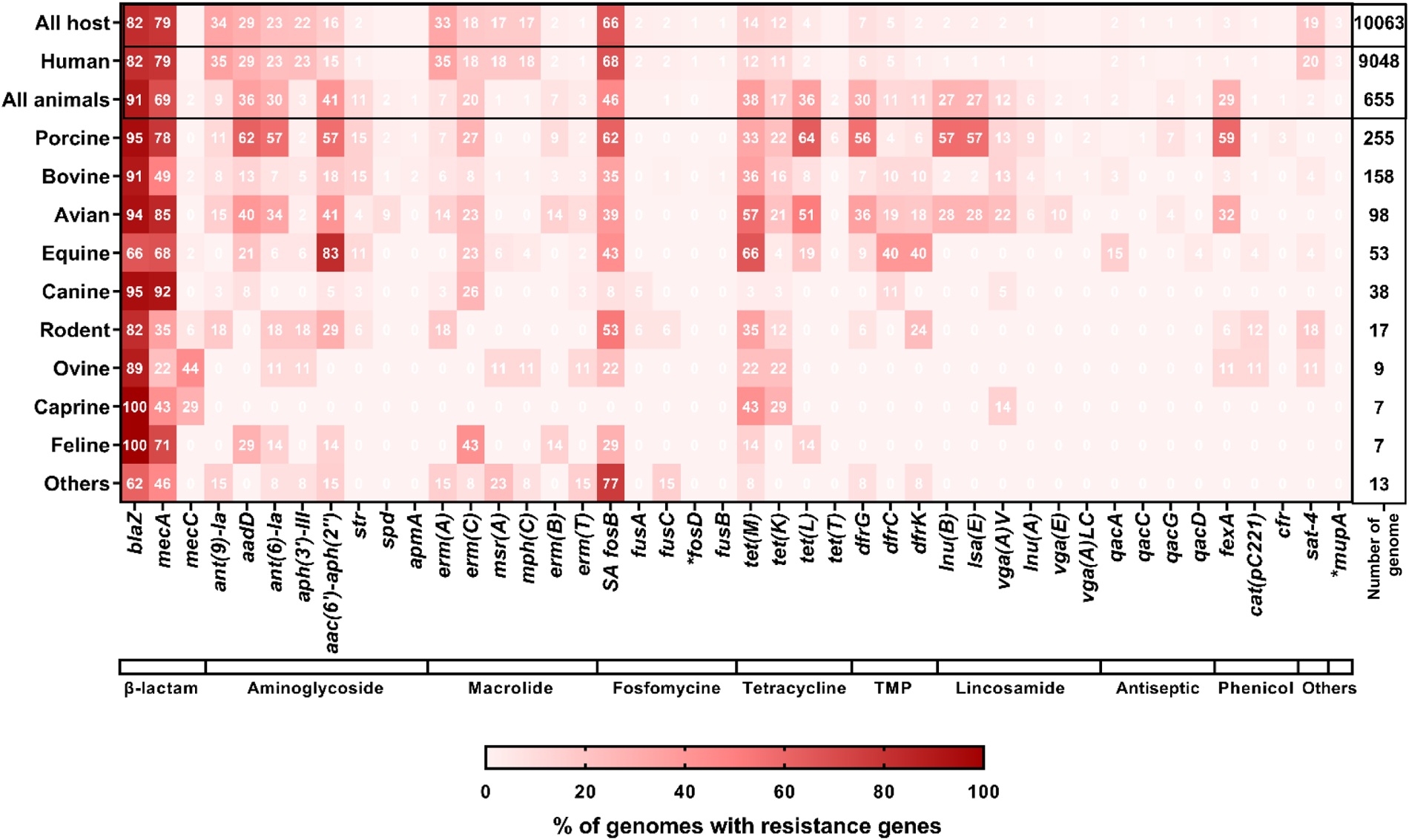
Occurrence of antibiotic resistance genes (ARGs) in *S. aureus* genomes. *S. aureus* hosts were indicated on the left of the Heatmap, detected ARGs on the bottom and the total number of genomes analyzed for each host on the right. Hosts grouped under the label ‘others’ correspond to dolphins (n=4), bats (n=4), fishes (n=2), cervid (n=1), shrimp (n=1) and monkey (n=1). The bar at the bottom of the Heatmap corresponds to the families of antibiotics with TMP for trimethoprim and ‘others’ for nucleoside (*sat-4*) and mupirocin (*mupA)* families. The percentages of genomes containing ARGs were obtained by dividing the number of genomes possessing the resistance gene by the total number of genomes on the top right of the Heatmap. These percentages are represented by different shades of red color, as shown at the bottom of the Heatmap. Only ARGs with a frequency >0.1% in Hm-SA and/or in An-SA, were represented. Genes exclusively present in Hm-SA are indicated by an asterisk.

The *blaZ* and *mecA* genes, conferring resistance to beta-lactam antibiotics, were commonly found regardless of the source and often co-harbored (n=417 in An-SA and n=5,913 in Hm-SA) (Figures 4 & S3). The *aac(6’)-aph(2’’)* (aminoglycoside), *tet(M)* and *tet(L)* (tetracyclines), *dfrG* (trimethoprim)*, lnu(B) and lsa(E)* (lincosamides) and *fexA* (phenicol) genes were more prevalent (present in >25% of genomes) in An-SA, while the *ant(9)-Ia* and *aph(3’)-III* (aminoglycosides) and *erm(A)* (macrolides) genes were more common in Hm-SA (Figure 4). Of note, the *aac(6’)-aph(2’’)* gene was found in 83% of the equine *S. aureus*, while it was present in less than 60% of the genomes of other animal hosts (Figure 4). Among animals, pig isolates presented the largest number of resistance genes, with 11 genes belonging to seven different families present in more than 40% of the genomes (Figure 4). In contrast, in bovine, only *blaZ* and *mecA* were present in more than 40% of the genomes (Figure 4). Certain ARGs exhibited restricted distribution patterns, as *vga(E)* present only in human-derived ST398 *S. aureus* strains or *vga(A)LC* and *apmA* identified in animal-derived ST398/ST9 strains. The ARG/ST associations were source-specific, except for *str*, *spd*, *ampA*, *erm(T)*, *dfrK* and *vga(E)*, which were predominantly (>80%) identified in ST398 genomes in both Hm-SA and An-SA (Figures S2C & D).

### Association of mobile genetic elements with antibiotic resistance genes in *S. aureus*

A Pearson correlation analysis revealed astrong positive correlation between the number of ARGs and the number of MGEs (excluding IS) in *S. aureus* genomes (r=0.57, p<0.0001, r=0.39, p<0.0001, for Hm-SA and An-SA, respectively). This correlation was particularly high for transposons, with R-values of 0.45 (p<0.0001) for Hm-SA and 0.60 (p<0.0001) for An-SA strains, but also for SCC elements (r=0.45, p<0.0001; r=0.48, p<0.0001) and plasmids (r=0.24, p<0.0001; r=0.46, p<0.0001).

The majority of ARGs (78% in Hm-SA, 75% in An-SA) were associated with MGEs (Figure5, Table S2). Plasmids (47% in An-SA and 42% in Hm-SA) were the predominant MGE harboring ARGs, followed by SCC elements (22% in Hm-SA and 11% in An-SA) and transposons (18% in Hm-SA and 12% in An-SA) (Figure 5, Table S2). The diversity of MGE/ARG co-occurrences was high since 21 and 12 different combinations of MGE/ARG were identified in Hm-SA and An-SA, respectively (Figure 5). The *spd*, *cat(pC221)*, *qacG*, *str*, *tet(T)* and *vga(A)LC* genes were exclusively found on plasmids, while *ant(9)-Ia*, *erm(A)*, *fexA*, *vga(A)V* and *vga(E)* were primarily associated with transposons, regardless of the *S. aureus* source (Figure 5). Several genes were associated with various types of MGEs, like *erm(B)* and *dfrK* which were frequently carried by plasmids or transposons and by putative plasmid/transposon tandem (Figure 5). *S. aureus* MGEs could be nested within each other, like plasmids or ICEs harboring transposons, or SCC*mec* elements hosting a diverse array of other MGEs (Figure 5).

**Figure 5.**
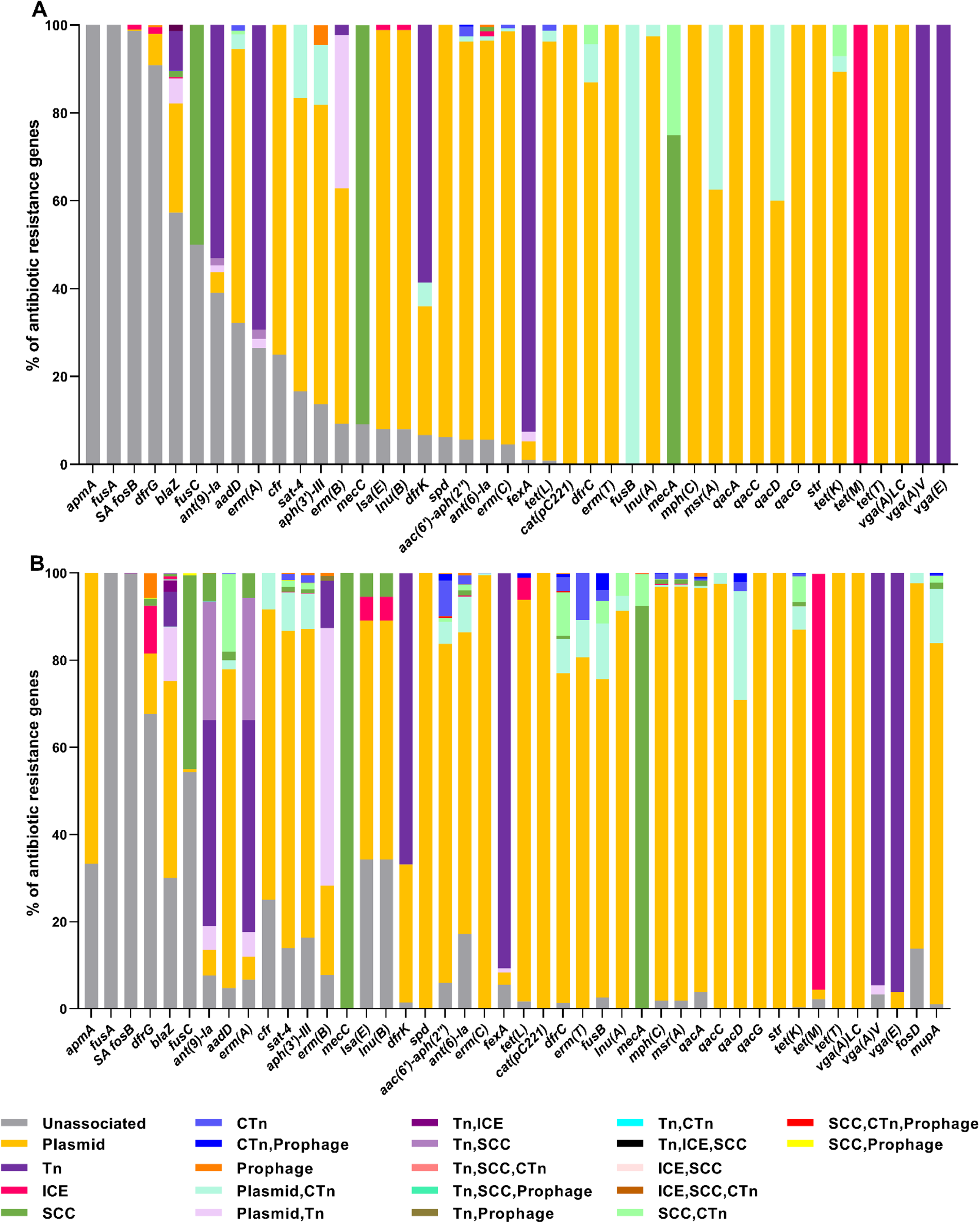
Associations of antibiotic resistance genes (ARGs) and mobile genetic elements (MGEs) in *S. aureus* genomes of animal (An-SA) (A) and human (Hm-SA) sources (B). Bar chart representing the percentage of associations between ARGs and MGEs, with each association assigned a color and indicated in the legend. When an ARG is not associated, the percentage is represented in gray. Only the ARGs with a frequency >0.1% in Hm-SA and/or An-SA, were depicted. The percentage of ARG corresponds to the number of ARG associated with MGE or a combination of MGEs divided by the total number of this ARG, identified in An-SA or Hm-SA. Tn: transposon; ICE: Integrative and Conjugative Element; SCC: Staphylococcal chromosomal cassette; CTn: composite transposon.

#### Plasmid/ARG associations

Each ARG was primarily associated with a specific Rep family (Figure 6). Rep1 and RepA_N were the dominant plasmid families associated with ARGs, either alone or in combination with other Rep (80%, 496/619 in Hm-SA; 78%, 164/210 in An-SA) (Figure 6). Within these, Rep1/*aadD* (8%, n=1,603/20,299) in Hm-SA and Rep1/*tet(L*) (6.5%, n=133/2,038) in An-SA, emerged as the most common. An-SAdisplayed two prevalentplasmid/multi-ARGs co-occurrences (Figure 7A).The first involved seven ARGs *aac(6’)-aph(2’’)*, *aadD*, *ant(6)-Ia*, *erm(C)*, *lnu(B)*, *lsa(E)* and *tet(L)* carried by a Rep1 plasmid (n=42). The second involved the *aadD*, *tet(L)* and *tet(T)* genes carried by an Inc18/Rep1 plasmid (n=13). In Hm-SA, the two most widespread associations were Inc18/Rep1/RepA_N (n=401) and Inc18/RepA_N (n=363), each associated with the same six ARGs *ant(6)-Ia*, *aph(3’)-III*, *blaZ, mph(C), msr(A) and sat-4* (Figure 7B).

**Figure 6.**
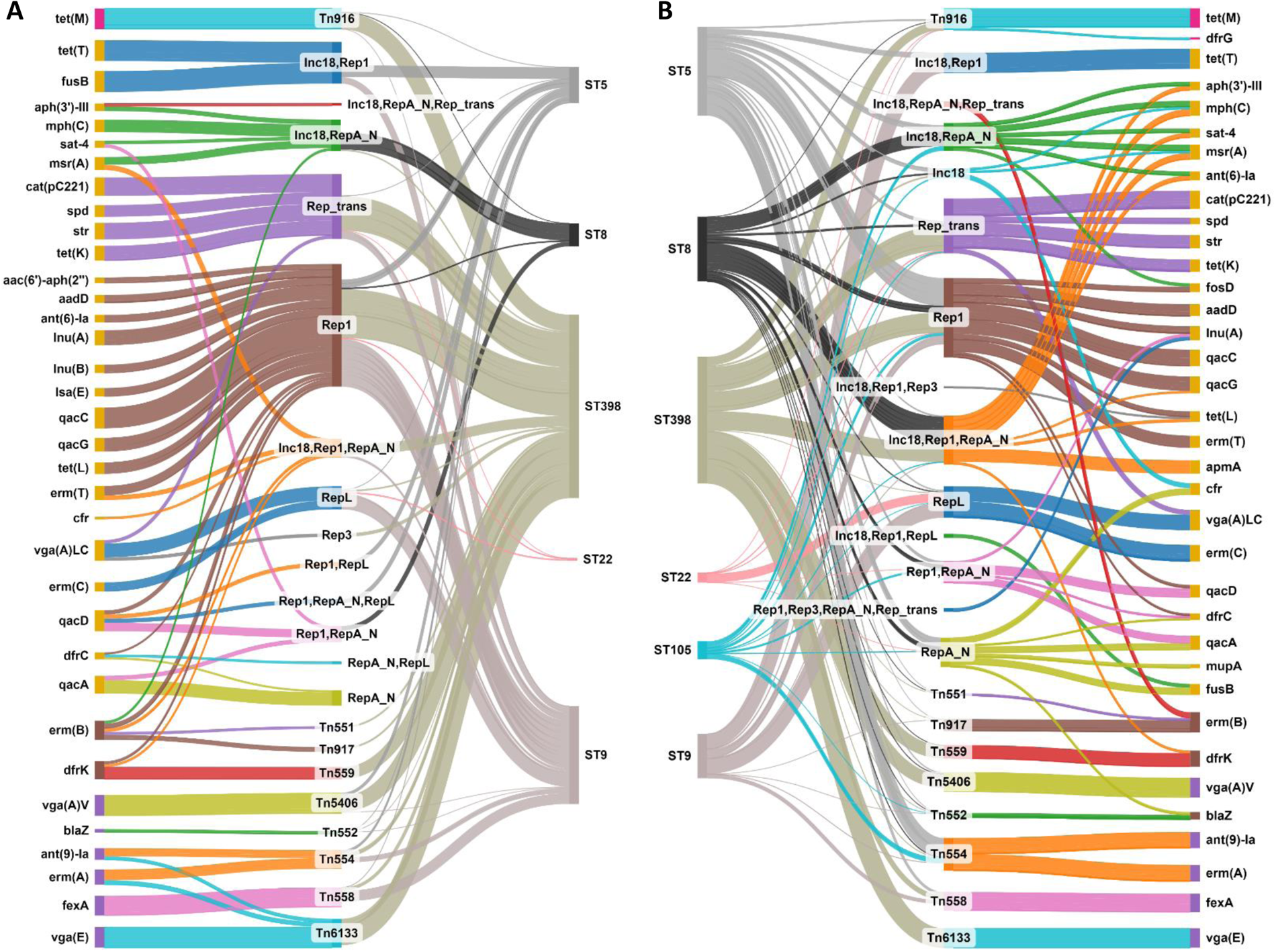
Co-occurrences of antibiotic resistance genes (ARGs) with plasmids, transposons and integrative and conjugative elements (ICEs) in *S. aureus* genomes of animal (An-SA) (A) and human (Hm-SA) (B) sources depending on sequence type (ST). Each ARG, MGE and ST were representedby a node that is linked by a flow when associated. The color of the ARG nodes corresponds to the type of MGE with which it is associated: pink for ICE, yellow for plasmid, purple for transposon and brown for plasmid and transposon. The thickness of the flowis proportional to the percentage of ARG associated with the MGE and the percentage of MGE associated with ST. This percentage is calculated by dividing the number of ARG associated with the MGE by the total number of these ARGs identified in An-SA or Hm-SA. Only ARGs with a frequency >0.1% in Hm-SA and/or in An-SA, and association frequencies of MGE/ARG >10% were represented. For the MGE/ST association, only the 6 most abundant STs in both Hm-SA and/or in An-SA were represented. The maximum thickness of the flow corresponds to 100%, as demonstrated by Tn*916*/*tet(M)* for (A) and Inc18,Rep1/*tet(T)* for (B). Sankey diagrams were created using SankeyMATIC (http://sankeymatic.com).

**Figure 7.**
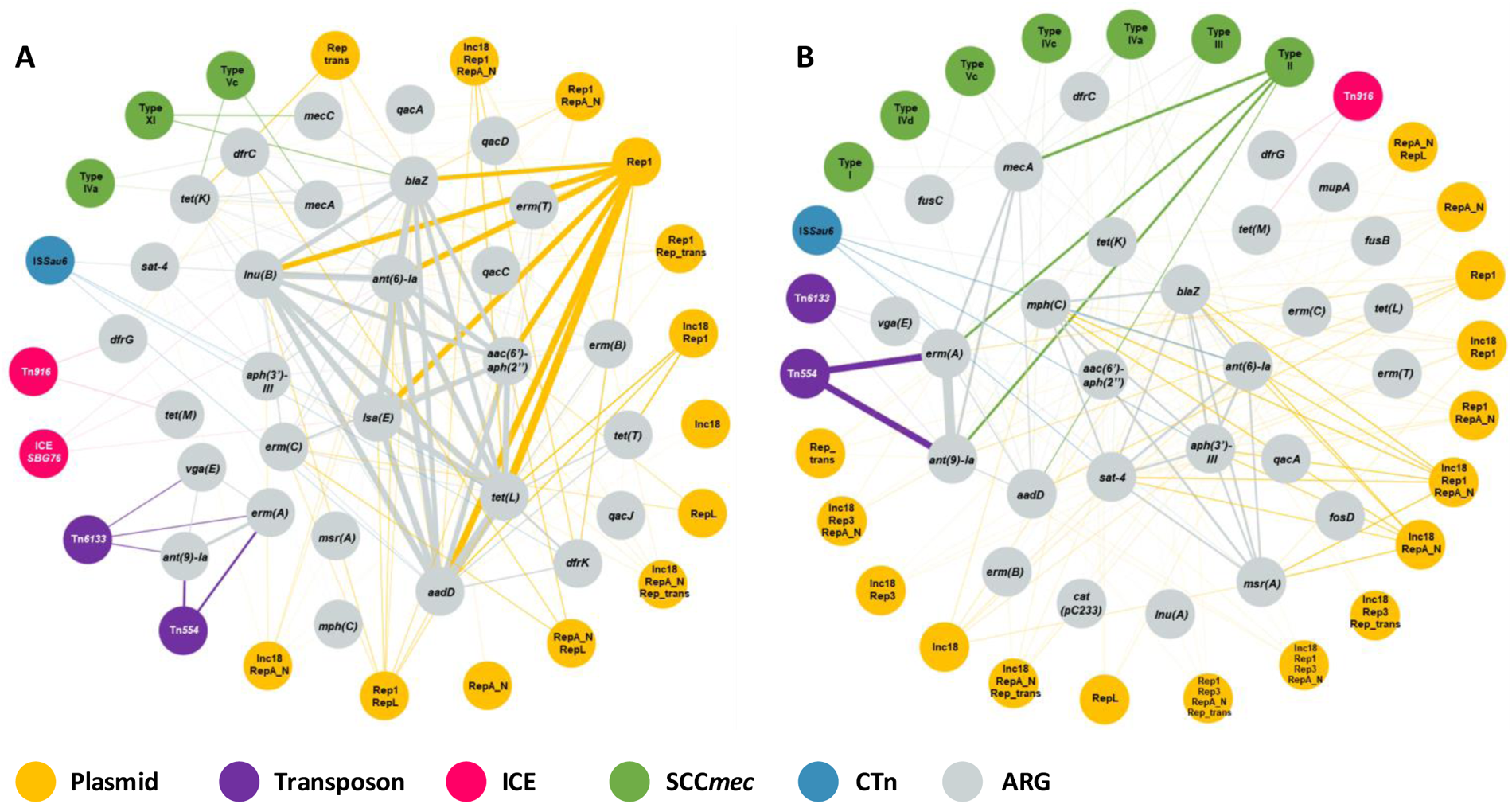
Co-occurrences between multiple antibiotic resistance genes (ARGs) and mobile genetic elements (MGEs) in *S. aureus* of animal (An-SA) (A) and human (Hm-SA) (B) sources. Associations were represented by links that indicate that ARGs were found within the same genome on the same MGE. The thickness of the line is proportional to the number of genomes carrying this co-occurrence. The thickest lines correspond to Rep1/*aac(6’)-aph(2’’)/aadD/ant*(*6*)*-Ia/erm(C)/lnu(B)/lsa(E)/tet(L)* association identified in 42 An-SA genomes and Tn*554*/*ant(9)-Ia*/*erm(A)* co-occurrence in 3,643 Hm-SA genomes. Only associations found in at least two An-SA genomes, or 10 Hm-SA genomes were shown. SCC: Staphylococcal chromosomal cassette; CTn: Composite transposon; ICE: Integrative and Conjugative Element.

#### Transposon/ARG associations

Each ARG was associated with one type of transposon and identical transposon/ARG associations were identified in both human and animal genomes (Figure 6). The Tn*554/ant(9)-Ia*, Tn*554*/*erm(A)* and Tn*558*/*fexA* associations were the most frequently detected in Hm-SA and An-SA, respectively(Figure 6). Apart from *erm(A)* and *ant(9)-Ia* found on Tn*554* and Tn*6133* or *erm(B)* associated with Tn*551* and Tn*917* (Figure 6), ARGs were found only on one type of transposon. Some ARGs, such as *blaZ*, were also found associated with both transposons and ICEs (Table S2). A strong co-occurrence between certain genes and transposons was observed, as for *lnu(G)*, *vga(A)V*, *vga(E)* or *fexA* genes that were predominantly associated with transposons (Figure 6). Transposon also harbored multiple ARGs. Notably, the Tn*554*/*erm(A)*/*ant(9)-Ia* association emerged as the most frequent MGE/multi-ARGs association in Hm-SA (n=3,643) and the second most common in An-SA (n=25) (Figure 7). Additionally, the Tn*6133*/*erm(A)*/*ant(9)-Ia*/*vga(E)* co-occurrence was identified in both Hm-SA (n=25) and An-SA (n=11) genomes (Figure 7).

#### Integrative and conjugative element/ARG associations

Overall, a minority (<9% of MGE/ARG associations) of ARGs were associated with ICEs, but associations were diverse (Table S2). The most widespread element was Tn*916*/*tet(M)* (Figure 6). This gene was itself usually located on ICEs (96%/100% in Hm-SA/An-SA) (Figure 6) and sometimes associated with other genes like *dfrG* (Figure 7).

#### Staphylococcal chromosomal cassette/ARG associations

The *mecA* gene was carried by several types of SCC*mec* cassettes, while *mecC* was exclusively associated with the type XI(8E) element. In addition to *mec* genes, 23 different ARGs were found on SCC*mec* elements and formed 97 SCC*mec*/ARG associations in Hm-SA genomes, while nine different ARGs were found forming 12 different SCC*mec*/ARG associations in An-SA strains (Table S2). In Hm-SA, the most common genes were *ant(9)-Ia* and *erm(A)* genes carried by a type II(2A) SCC*mec* element (1,513 for *ant(9)-Ia* and 1,473 for *erm(A)*) and in An-SA genomes *blaZ* was always associated with type XI(8E) cassette carrying *mecC* (n=10) (Table S2). The SCC*mec*/multi-ARGs associations, like type Vc/*mecA*/*tet(K)* and type IVa/*mecA*/*dfrC*, were found in both humans (n=27 and and animals even though their proportions differed depending on the source (n=27 and 41 in Hm-SA, n=8 and 3 in An-SA) (Figure 7, Table S2).

#### Prophage/ARG associations

ARGs were rarely associated with prophages (0.3% in Hm-SA, 0.1% An-SA genomes). In Hm-SA genomes, the most common prophage/ARG associations were phi2958PVL/*dfrG* (n=27) and SPbeta_like/*aac(6’)-aph(2’’)* (n=23) (Table S2). This prophage was found associated with 17 different ARGs (Table S2). Prophage/ARG occurrences were not exclusive, i.e. *blaZ* could be associated with 10 different types of prophages. In An-SA genomes, only two associations were identified, i.e. *aac(6’)-aph(2’’)*, *aph(3’)-III* and *ant(6)-Ia* carried by a SPbeta-like prophage, and *dfrG* associated with YMC/09/04/R1988.

#### Composite transposon/ARG associations

These MGEs were only associated with about 5% of ARGs in both Hm-SA and An-SA (Table S2). IS*Sau6* was the most common CTn associated with several ARGs (24 different ARGs in Hm-SA and 15 in An-SA genomes) (Table S2). The most frequently identified co-occurrences were IS*Sau6*/*mecA*, harbored by a SCC*mec* element, in An-SA strains (n=103), but also abundant in Hm-SA (n=534), even if IS*Sau6*/*aadD*, also located on SCC*mec*, was most widespread in Hm-SA (n=547).

## Discussion

Although MGEs are recognized key players in the dissemination of AMR, the association between MGEs and ARGs has been understudied in Gram-positive bacteria, particularly in *S. aureus*. While previous studies focused either on MGEs and/or ARGs (8, 9, 18, 19), our large-scale analysis of over 10,000 publicly available *S. aureus* genomes from both animal and human sources is, to our knowledge, the first to reveal the intricate interplay between mobilome and resistome. Our analysis highlighted a remarkable diversity of STs (n=433), MGEs (n=278) and ARGs (n=78) in *S. aureus,* as well as the routes of their dissemination within the species. Our study, like all studies based on publicly available genomes, has some limitations imposed by the bias in sequenced genomes deposited in the NCBI database. Due to sequencing costs, only strains harboring the most critical resistance genes for human and animal health were analyzed. This resulted in an over-representation of MRSA strains (80% Hm-SA and 71% An-SA) and the under-representation of An-SA, except for *mecA*-positive ST398 and ST9 isolates of porcine origin (n=143/655 An-SA, 22%) and MRSA of bovine-origin (n=81/655 An-SA, 12%). Additionally, our analyses included both complete and draft genomes, which may have resulted in the exclusion of elements spanning several contigs. However, since our study aimed to capture the diversity of MGEs and ARGs rather than quantify them, this bias did not significantly interfere with our analyses.

Our study showed a wide distribution and a large diversity of ARGs in *S. aureus* of human and animal origin, even larger than the one reported in the review by Schwartz *et al.* (7), and capable of conferring resistance to all antibiotic families known to treat staphylococcal infections. The most widespread genes were *blaZ* and *mecA*, as also pointed out in a recent study of AMR in publicly available *S. aureus* genomes (20). A recent study on the resistome/mobilome association in ESKAPE pathogens highlighted the overall high prevalence of genes conferring resistance to aminoglycosides, chloramphenicol, trimethoprim and tetracyclines (21). Specific genes conferring resistance to these antibiotic families were also particularly frequent in our study dedicated to *S. aureus*, and their wide dissemination is consistent with antibiotic use in both human and veterinary medicine. Globally, our study revealed a higher diversity of ARGs in Hm-SA and a larger set of human-specific genes. This could be explained by the larger number of genomes analyzed, but also by the higher exposure of humans to antibiotic selection in various environments (hospitals, farm environments or food). Only one gene, *optrA*, was specific to An-SA. This gene, first discovered inhuman and animal enterococci on a conjugative plasmid (22), and then observed in *Staphylococcus sciuri* of porcine origin (23), is conferring resistance to phenicols. It was found on *S. aureus* chromosome near the SCC*mec*, but also on a plasmid with the *cfr* gene (23). In our study, *optrA* was identified on the chromosome and not associated with an MGE: nevertheless, it was located <10kb from the *fexA* gene carried by Tn*558*, an MGE/ARG association already found on the plasmid carrying *cfr* in other staphylococci (24, 25). Besides these host specificities, a large set of genes, among which *aph(3’)-III*, *dfrC*, *fusA*, *tet(T)*, *apmA* and *vga(E)*, were shared between An-SA and Hm-SA. This suggests that numerous ARGs have no host boundaries and can be disseminated from human *S. aureus* to animals and vice versa.

The *S. aureus* mobilome was largely composed of IS and phages, which rarely carried ARGs, followed by plasmids and transposons that were identified as the main carriers of ARGs for this species. Our study also highlighted that plasmids exhibited the highest diversity among MGE, with 297 different *rep* genes combinations often forming mosaic plasmids with multiple *rep* genes (37% in Hm-SA and 23% in An-SA) (12, 26). Previous studies mainly reported the association of only two *rep* genes, belonging to the same or two different plasmid families (27, 28). However, our work demonstrates that as many as eight different *rep* genes can be found in a single plasmid. Unlike plasmids circulating in Enterobacterales, those identified in *S. aureus* present few Rep proteins incompatibilities. This facilitates the formation of mosaic plasmids, enabling *S. aureus* to adapt more effectively to diverse hosts and environmental conditions (27, 29–31). Furthermore, the presence of numerous insertion sequences (IS elements) and the rolling-circle replication(RCR) system of small plasmids(<10 kb) favors the recombination and fusion of several small plasmids, resulting in unique large hybrids carrying multiple *rep* genes (27, 32).

The prevailing view for a long time held that *S. aureus* possessed very few plasmids, and horizontal gene transfer was believed to occur primarily through transduction. However, studies on Gram-positive bacteria showed that plasmids can be mobilized by using the conjugation system of other plasmids or ICEs present in the strain (33–35). A study in *Bacillus subtilis* showed that plasmids can be mobilized by the *ori*T of ICE*Bs1,* an ICE of the Tn*916* family (34, 35). In *S. aureus*, plasmids can be mobilized if they possess a relaxase or by forming co-integrates that are subsequently resolved by “conduction” (36, 37). O’Brien *et al.* (38) showed that the conjugative plasmid pWBG749 could mobilize the transfer of numerous small plasmids carrying an *ori*T similar to its own (38, 39). Thus, plasmids carrying ARGs, even non-conjugative ones, could be easily transferred between cells, highlighting their central role in the dissemination of these genes (15).

Often misclassified in the literature as transposons or plasmids (40), ICEs are also capable of HGT. Studies on enterococci and streptococci showed that these MGEs play a crucial role in the dissemination of ARGs (41). The well-known Tn*916* family was found in many Firmicutes, always carrying the *tet(M)* gene (42–44). This element exhibits a similar backbone sequence between *S. aureus* and streptococci (43). Within *S. aureus*, it was first described in the Mu50 strain and named Tn*5801* (45). Our study identified another ICE, ICE*6013*, as the most prevalent family typically harboring few resistance genes itself. However, it can carry other MGEs like Tn*552*, often associated with the *blaZ gene*. This association of multiple MGEs with ARG was first identified in *S. aureus* ST239 (14). Despite sharing some protein similarities with the ICE*Bs1* of *Bacillus subtilis*, ICE*6013* belongs to a distinct family (14). Other MGEs were also found nested within each other, like transposons found within plasmids and SCC*mec* cassettes. For instance, Tn*554*/*ant*(*9*)*-Ia*/*erm(A)* was found on both Rep1 plasmids and SCC*mec* type II(2A) cassettes. The interweaving of MGEs significantly amplifies the potential for dissemination of ARGs (14, 40).

Beyond revealing the remarkable diversity of ARGs and MGEs in *S. aureus*, our study importantly evidenced numerous associations between these elements. The high degree of similarities between MGE/ARG associations observed in An-SA and Hm-SA strongly suggested unrestricted dissemination of MGEs across hosts. This finding implies that An-SA could serve as a reservoir for Hm-SA and vice versa. Our results identified a few dominant STs within Hm-SA or An-SA populations, typically harboring the same set of ARG/MGE associations. This observation highlights clonal expansion as a major driver of dissemination in *S. aureus*. This is the case for *S. aureus* ST22, initially identified in United Kingdom hospitals (46), which has become a globally disseminated epidemic clone and has spread across Europe and worldwide, impacting both healthcare settings and the community, including domestic animals like dogs and cats (47, 48). Due to its pandemic character, very few variations were observed in ST22 MGE content in genomes available in public databases. Although our study did not capture evidence of such events, the reported presence of a plasmid carrying via Tn*558* the *cfr* and *fexA* genes in Irish ST22 isolates (25) demonstrates the potential for horizontal transfer even within this otherwise stable clonal lineage. Besides this specific example of a rare event, our results also evidenced more frequent transfers. Notably,within the same ST, the observed diversity of MGEs and ARGs demonstrates that the mobilome and resistome are sources of significant genomic plasticity, despite core genome similarity. Furthermore, our analysis revealed an intense spread of certain MGE/ARG associations across diverse STs (Figure 6). For instance, the Rep1/*aadD* association was found in all five most prevalent STs. Additionally, the Tn*916*/*tet(M)* is primarily associated with ST398 in An-SA (80% of identified *tet(M)* genes are in ST398), while in Hm-SA, it was found in both ST398 (40%) and ST239 (20%). This suggests potential host adaptation, in line with recent studies demonstrating the role of *tet(M)*-carrying Tn*916* transposon, facilitating the jump from An-SA to Hm-SA (49). Despite lacking their own transfer systems and residing primarily within chromosomes, transposons also contribute to *S. aureus* genome plasticity and resistome dissemination among STs.

Our findings revealed the presence of transposons in diverse STs, like Tn*554*/*ant*(*9*)*-Ia*/*erm(A)* which was identified in more than 12 STs, and Tn*558*/*fexA* found in 10 An-SA STs. This widespread distribution across different epidemic clones highlights the horizontal propagation of these elements, suggesting their important role in shaping the resistome and, more broadly the genomes of *S. aureus*.

In conclusion, this study revealed the remarkable abundance and diversity of MGEs and ARGs within *S. aureus* genomes, primarily associated with plasmids and transposons. While it is known that *S. aureus* disseminate through clonal waves, our results showed that these clones can also evolve through horizontal acquisition of MGE/ARG associations that are promoting the dissemination of resistance genes. This study provides additional evidence that An-SA can serve as reservoirs for Hm-SA and vice versa, and that the mobilome’s only limitation is its transfer efficiency. Plasmids in *S. aureus*, as primary reservoirs of ARGs, are key players of AMR dissemination. This highlights the critical need to elucidate the mechanisms governing the epidemic success of MGEs, particularly those implicated in ARG transfer.

## Materials and methods

### Genome characterization and phylogenetic analysis

Genome assemblies (complete and draft genomes) of *S. aureus* of animal (n=1,436) and human sources (n=11,888) were downloaded from the NCBI public database in October 2021. Quality and completeness of assemblies were evaluated using QUAST v.5.0.2 (50) and BUSCO v.5.2.1 (51) (Figure S4). All contigs <1000 bp were removed from the analysis. Genomes were characterized according to their ST using MLST v.2.1.1 (52). Transposons, ICE, IS and CTn were identified using MobileElementFinder v.1.0.3 (80% coverage and 70% identity) (53). Plasmid replicons (*rep* coding gene) were detected using PlasmidFinder v.2.1.6 software (databases: v.2021-11-29) (60% coverage and 90% identity) (54, 55). SCCs were identified using SCC*mec*Finder v.1.2.1 (database extended, 60% coverage and 90% identity) (54, 56, 57). ARGs were identified using ResFinder v.4.1.7 (database: v.2021-09-22) (60% coverage and 80% identity) (54, 58) and RGI-CARD (Comprehensive Antibiotic Resistance Database) (59) (percentage length of the reference sequence >200). Co-occurrence was considered when several ARGs were identified within the same strain. A phylogenetic tree was constructed using pyMLST v.2.1.3 with default parameters (60). A neighbor-joining tree based on the *S. aureus* core genome was constructed using cgMLST software (https://www.cgmlst.org/ncs) (61). To avoid duplicates, only one genome with identical MGE and ARG content, geographical origin, collection year and NCBI bioProject was selected in each group determined by cgMLST with a maximum distance of 10 different alleles (Figure S4). Following this selection, a total of 9,408 genomes of Hm-SA and 655 genomes of An-SA, carrying at least one ARG and one MGE (except IS) were included in this study.

### Identification and characterization of detected mobile genetic elements

#### Plasmids identification

PlasForest (54, 62, 63), RFPlasmid for *Staphylococcus* species (64), Plasmer (65) and Plasflow (66) were used with default parameters to identify whether contigs were plasmidic or chromosomal. A contig was considered plasmidic if predicted by at least three software. Due to the fragmented nature of draft genomes, some plasmids were split into multiple contigs during assembly. To identify contigs belonging to the same plasmid, those identified as plasmidic were analyzed using BLASTN. Contigs were considered part of the same plasmid if they shared more than 60% identity and coverage with the same reference plasmid sequence. Some previously unclassified contigs were identified as plasmidic due to their similarity to known plasmids. Plasmids were classified according to the active protein domain of the Rep protein that was encoded by *rep* gene. All *rep* genes identified on chromosomal contigs were excluded from the analysis.

#### Composite transposons identification

A CTn was predicted when two IS were at a specified distance from each other. The threshold value was 12kb according to the length of the largest CTn identified in *S. aureus* (8).

#### Staphylococcal Chromosomal Cassettes identification

Boundaries of SCC was fixed with *orfX* gene and the end was adjusted based on the size of the cassette type (https://www.sccmec.org/). SCCs found scattered across multiple contigs were adjusted based on gene order (synteny) to define their limits. SCC*mec*Finder v.1.2.1 did not determine the cassette type for some SCC elements, considered as “indeterminate”. SCC lacking *mecA* or *mecC* genes were classified “SCC” instead of SCC*mec*.

#### Prophages identification

These elements were identified using PHASTEST, last accessed October 2023 (67–69). Only prophages with a “complete” score were retained. Prophages were named according to the bacteriophage exhibiting the closest similarity, determined by both sequence identity and the number of common genes.

#### Integrative and Conjugative Elements identification

In addition to MobileElementFinder v.1.0.3 detection, a BLASTP analysis was performed against a custom database created from literature and ICEBerg 2.0 database (15, 45, 70, 71). This database included conserved components of the two main ICE families, Tn*916* and ICE*6013*, identified basedon a previous study (70) (Figure S5A). Only conserved elements meeting the following criteria were considered as ICEs: proximity between elements, number of conserved elements on the same contig and integrity of the gene (Figure S5B). ICEs boundaries were extended by 10kb at each end of the ICE, reflecting the average size of Staphylococci ICE (approximately 30kb).

### Identification of associations between antibiotic resistance genes and mobile genetic elements

To assess the correlation between the number of ARGs and the number of each MGE within a genome, the Pearson correlation coefficient was calculated using GraphPad Prism 9.5.1 (GraphPad Software Inc., CA, USA). MGE/ARG associations were identified through co-localization analysis when the MGE region entirely overlapped the ARG region. This rule was applied only for transposons, ICE, SCC, prophages and CTn. For plasmids, an ARG was considered associated when it was localized on a plasmidic contig (see above). For plasmidic contig lacking *rep* gene, associated ARGs were considered carried by a plasmid of non-determinate (ND) type. Unclassified contigs identified as associated with SCC, prophage, ICE were reclassified as chromosomal. When multiple ARGs were found within the same MGE from a single strain, the MGE was considered to harbor multiple ARGs.

## Supporting information

Supplementary figures with legends

## Acknowledgments

This work was supported by PhD grant from INRAE (French National Research Institute for Agriculture, Food and the Environment) and ANSES (French National Agency for Food, Environmental and Occupational Health and Safety). We are grateful to the Genotoul bioinformatics platform Toulouse Midi-Pyrenees, France for providing support and storage resources. We address a special thanks to Marie-Stephane Trotard from Genotoul, for his expertise, advice and installation of all bioinformatics software. We thank Gaëtan Durand for the python scripts that enabled us to analyze a large volume of data.

