## Supplementary figures with legends for "The interplay between Mobilome and Resistome in *Staphylococcus aureus*"

### Supplementary data

**Table S1. Database of MGEs and ARGs detected in *S. aureus* of human and animal sources.**

**Table S2. Database of MGE/ARG associations in *S. aureus* of human and animal sources.**

**Figure S1. Occurrence and diversity of sequence types (STs) in *S. aureus* in relation to their host (A) and depending on human (Hm-SA, black) and animal (An-SA, white) sources (B). (A)** The frequency of each ST corresponds to the number of strains classified by their respective ST and host, divided by the total number of strains of that ST. Each host corresponds to a color as defined on the right of the graph. Hosts grouped under the label 'others' correspond to dolphins (n=4), bats (n=4), fishes (n=2), cervid (n=1), shrimp (n=1) and monkey (n=1). Only STs with a frequency >1% in Hm-SA and/or An-SA, were represented. **(B)** The frequencies corresponded to the number of genomes of each type divided by the total number of genomes according to their source (Hm-SA n=9,408 and An-SA n=655).

**Figure S2. Frequency of Mobile genetic elements (MGEs) and antibiotic resistance genes (ARGs) depending on sequence types (STs) in *S. aureus* of animal (An-SA) (A&C) and human (Hm-SA) (B&D) sources.** The frequencies of ARGs and MGEs are calculated based on the number of genomes carrying each ARG or MGE within their respective ST, divided by the total number of strains carrying each ARG or MGE. Only ARGs with a frequency >0.1% in Hm-SA and/or in An-SA, were represented.

**Figure S3. Association between antibiotic resistance genes (ARGs) in *S. aureus* of animal (An-SA) (A) and human (Hm-SA) (B) sources.** Links indicated that the two genes were found within the same genome. The thickness of the line was proportional to the amount of genome carrying this co-occurrence. The thickest lines correspond to *blaZ/mecA* association identified in 417 An-SA genomes and 5,913 Hm-SA genomes. Only ARG found in >1% of Hm-SA or An-SA genomes were represented. Each node is colored according to the ARG antibiotic family. 'TMP' corresponds to Trimethoprim, 'Others' to nucleoside (*sat-4)* and mupirocin (*mupA)* antibiotic families.

**Figure S4. Selection of *S. aureus* genome of human (Hm-SA) and animal sources (An-SA).** This decision tree shows the process that was followed to select the genomes used in our study. One genome with only a large amount of insertion sequence (n=87) (GCA_014876755.1) (a), and one with incoherent cgMLST profile (GCA_001921685.1) (b) were excluded. (c) Six genomes for which not all analyses were achievable with NCBI data, due to the unavailability of certain files, were also excluded (GCA_014337115.1; GCA_014336505.1; GCA_014353635.1; GCA_014353595.1; GCA_014353515.1; GCA_900018155.1). MGE: Mobile genetic element; ARG: antibiotic resistance genes IS: Insertion sequence; Tn: transposon; ICE: Integrative and Conjugative Element; SCC: Staphylococcal chromosomal cassette; CTn: composite transposon.

**Figure S5. Criteria for integrative and conjugative elements (ICEs) identification and validation after BLASTP analyses. (A)** List of proteins or protein domains used to identify the presence of one of the two ICE families. **(B)** Decision tree showing how the presence or absence of an ICE has been validated.

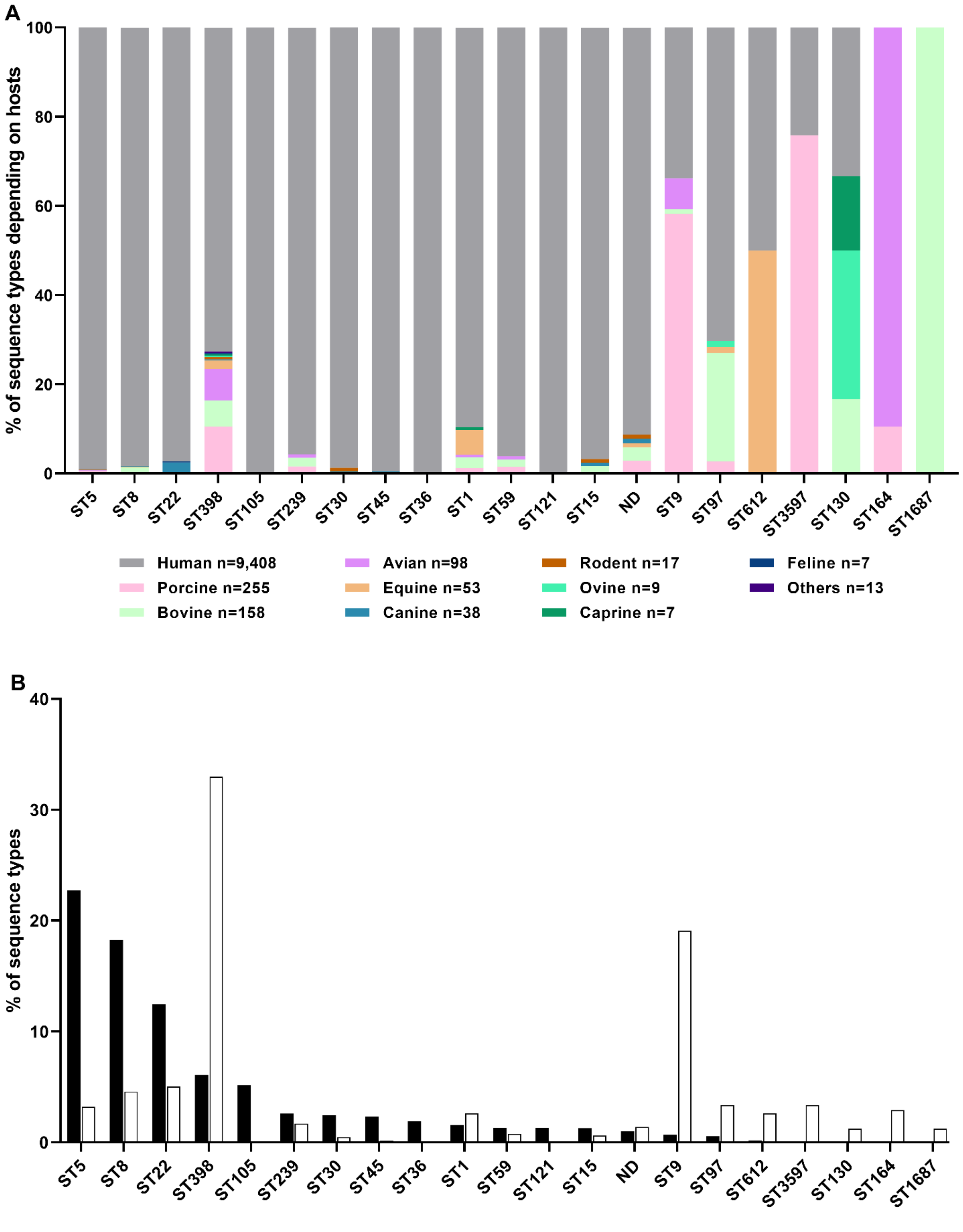

A
% of sequence types depending on hosts
100
80
60
40
20
0
ST5
ST8
ST22
ST398
ST105
ST239
ST30
ST45
ST36
ST1
ST59
ST121
ST15
ND
ST9
ST97
ST612
ST3597
ST130
ST164
ST1687
Human n=9,408
Porcine n=255
Bovine n=158
Avian n=98
Equine n=53
Canine n=38
Rodent n=17
Ovine n=9
Caprine n=7
Feline n=7
Others n=13
B
% of sequence types
40
30
20
10
0
ST5
ST8
ST22
ST398
ST105
ST239
ST30
ST45
ST36
ST1
ST59
ST121
ST15
ND
ST9
ST97
ST612
ST3597
ST130
ST164
ST1687


**Figure S1.**

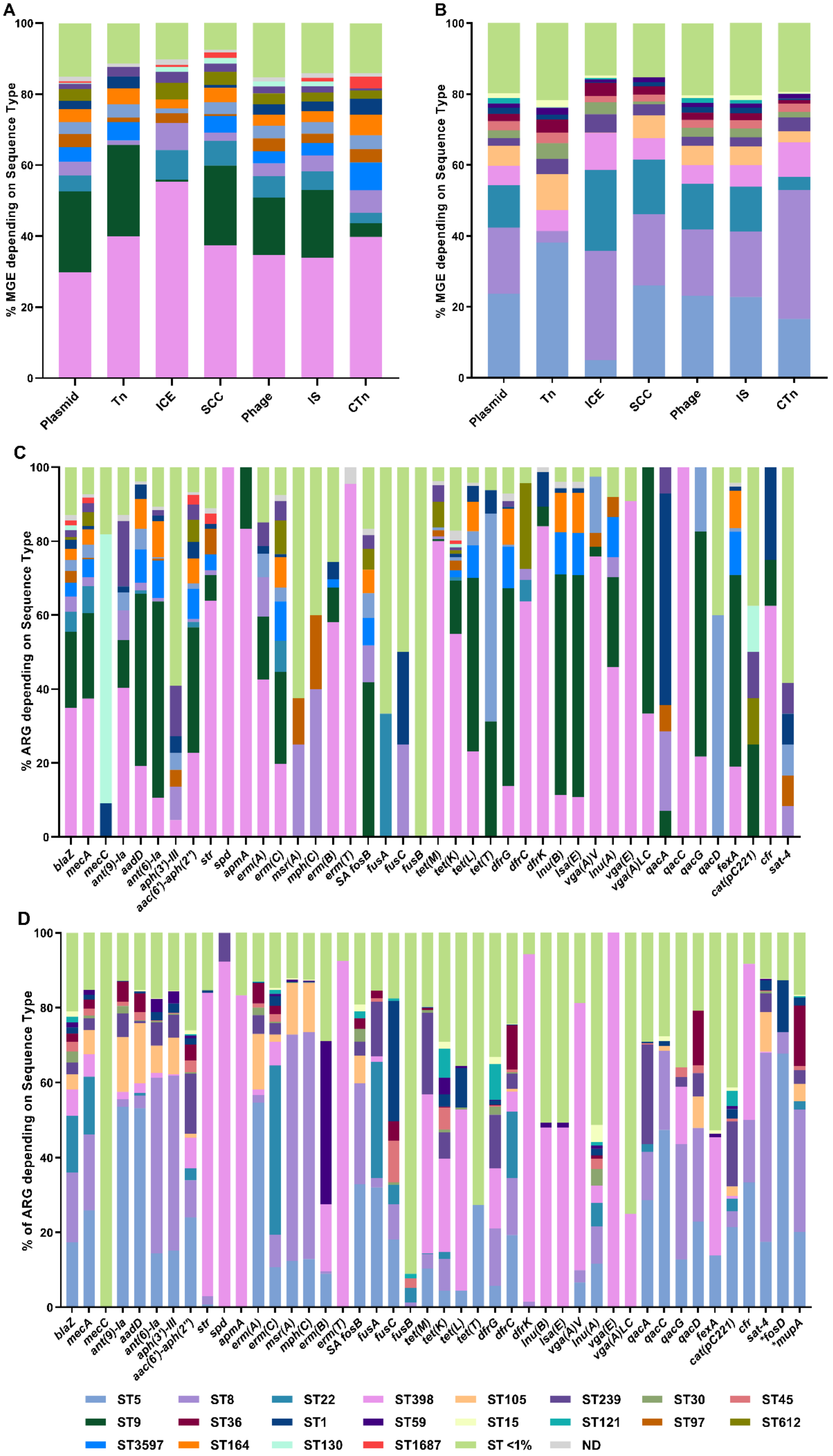

A
% MGE depending on Sequence Type
0
20
40
60
80
100
Plasmid
Tn
ICE
SCC
Phage
IS
CTn
B
% MGE depending on Sequence Type
C
% ARG depending on Sequence Type
blaZ
mecA
mecC
ant(9)-Ia
aadD
ant(6)-Ia
aph(3')-III
aac(6')-aph(2'')
str
spd
apmA
erm(A)
erm(C)
msr(A)
mph(C)
erm(B)
erm(T)
SA fosB
fusA
fusC
fusB
tet(M)
tet(K)
tet(L)
tet(T)
dfrG
dfrC
dfrK
lnu(B)
lsa(E)
vga(A)V
lnu(A)
vga(E)
vga(A)LC
qacA
qacC
qacG
qacD
fexA
cat(pC221)
cfr
sat-4
D
% of ARG depending on Sequence Type
*fosD
*mupA
ST5
ST8
ST22
ST398
ST105
ST239
ST30
ST45
ST9
ST36
ST1
ST59
ST15
ST121
ST97
ST612
ST3597
ST164
ST130
ST1687
ST <1%
ND


**Figure S2.**

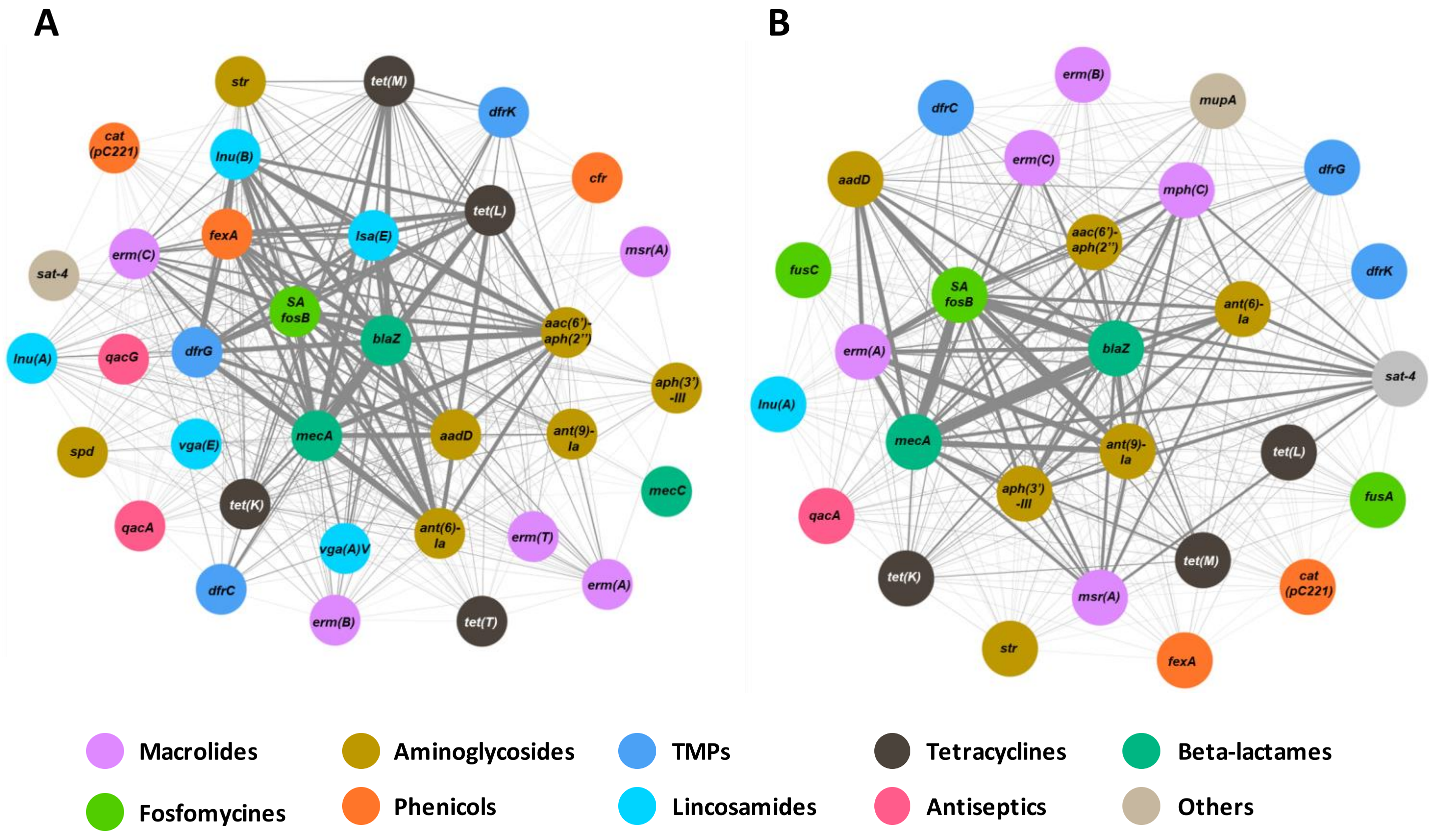
A
str
tet(M)
dfrK
cat (pC221)
lnu(B)
cfr
tet(L)
fexA
lsa(E)
msr(A)
erm(C)
sat-4
SA fosB
blaZ
aac(6')-aph(2'')
lnu(A)
qacG
dfrG
aph(3')-III
mecA
aadD
ant(9)-Ia
spd
vga(E)
mecC
tet(K)
qacA
vga(A)V
ant(6)-Ia
erm(T)
dfrC
erm(A)
erm(B)
tet(T)
B
erm(B)
dfrC
mupA
erm(C)
aadD
mph(C)
dfrG
aac(6')-aph(2'')
fusC
SA fosB
dfrK
ant(6)-Ia
blaZ
erm(A)
sat-4
lnu(A)
mecA
ant(9)-Ia
tet(L)
aph(3')-III
fusA
qacA
tet(M)
tet(K)
msr(A)
cat (pC221)
str
fexA
Macrolides
Aminoglycosides
TMPs
Tetracyclines
Beta-lactames
Fosfomycines
Phenicols
Lincosamides
Antiseptics
Others

**Figure S3.**

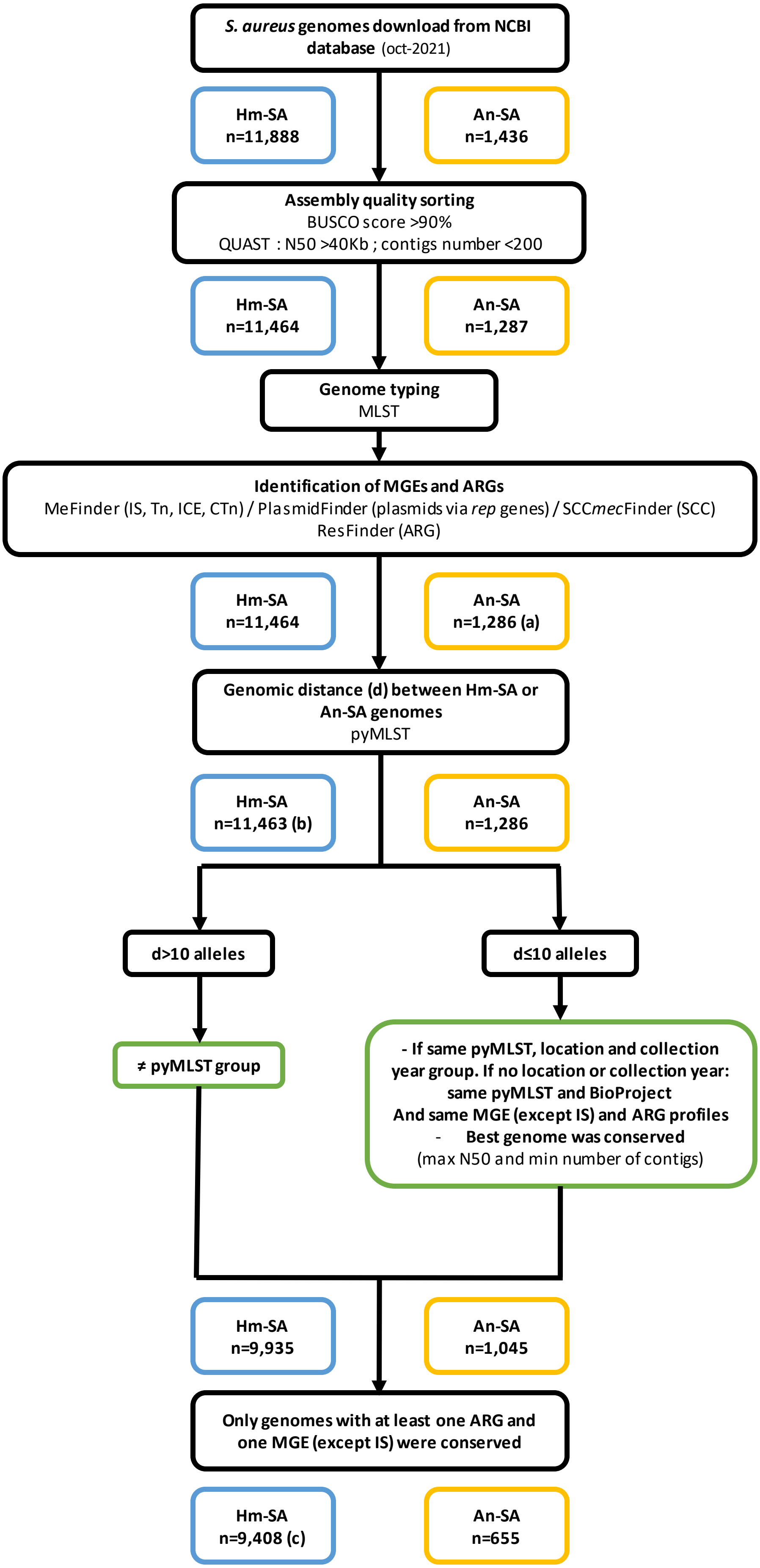
*S. aureus* genomes download from NCBI database (oct-2021)
Hm-SA n=11,888
An-SA n=1,436
Assembly quality sorting
BUSCO score >90%
QUAST : N50 >40Kb ; contigs number <200
Hm-SA n=11,464
An-SA n=1,287
Genome typing
MLST
Identification of MGEs and ARGs
MeFinder (IS, Tn, ICE, CTn) / PlasmidFinder (plasmids via *rep* genes) / SCC*mec*Finder (SCC)
ResFinder (ARG)
Hm-SA n=11,464
An-SA n=1,286 (a)
Genomic distance (d) between Hm-SA or An-SA genomes
pyMLST
Hm-SA n=11,463 (b)
An-SA n=1,286
d>10 alleles
d≤10 alleles
≠ pyMLST group
- If same pyMLST, location and collection year group. If no location or collection year: same pyMLST and BioProject
And same MGE (except IS) and ARG profiles
- Best genome was conserved
(max N50 and min number of contigs)
Hm-SA n=9,935
An-SA n=1,045
Only genomes with at least one ARG and one MGE (except IS) were conserved
Hm-SA n=9,408 (c)
An-SA n=655

**Figure S4.**

**A**

| ICE family | Protein Name | Protein Family | GeneBank accession number |
|---|---|---|---|
| ICE6013 | YddE | VirB4 domain protein | ACI48609.1 |
| | YdcQ | FtsK/SpoIIIE domain-containing protein | ACI48608.1 |
| | RstA | relaxase of the MobT family | ACI48613.1 |
| Tn916 | ORF16 | Conjugative transposon protein ATP-binding protein | CAQ49388.1 |
| | ORF21 | FtsK/SpoIIIE domain-containing protein | CAQ49393 |
| | ORF22 | Conjugative transposon protein YdcP family protein, DUF261 | CAQ49394.1 |

**B**

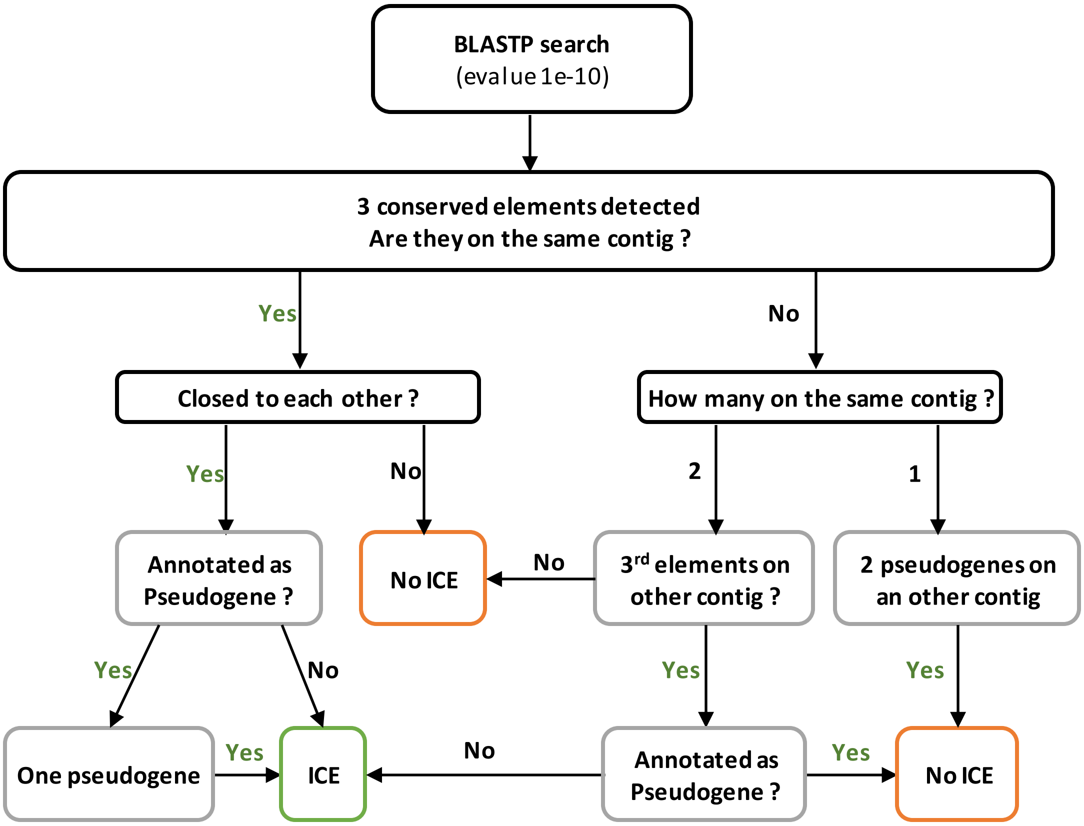


**Figure S5.**
